# Non-productive Binding Modes as a Prominent Feature of Aβ_1-40_ Fiber Elongation: Insights from Molecular Dynamics Simulation

**DOI:** 10.1101/287383

**Authors:** Rajiv K Kar, Jeffrey R Brender, Anirban Ghosh, Anirban Bhunia

## Abstract

Amyloid formation has been implicated in a number of neurodegenerative diseases. The elongation of amyloid fibers is thermodynamically strongly favorable but kinetic traps exist where the incoming monomer binds in an incompatible conformation that blocks further elongation. Unfortunately, this process is difficult to follow experimentally at the atomic level. It is also too complex to simulate in full detail and thus so far has been explored either through coarse-grained simulations, which may miss many important interactions, or full atomic simulations in which the incoming peptide is constrained to be near the ideal fiber geometry. Here we use an alternate approach starting from a docked complex in which the monomer is from an experimental NMR structure of one of the major conformations in the unbound ensemble, a largely unstructured peptide with the central hydrophobic region in a 3_10_ helix. A 1000 ns full atomic simulation in explicit solvent shows the formation of a metastable intermediate by sequential, concerted movements of both the fiber and monomer. A Markov state model shows the unfolded monomer is trapped at the end of the fiber in a set of interconverting anti-parallel β-hairpin conformations. The simulation here may serve as a model for the binding of other non-β-sheet conformations to amyloid fibers.

## Introduction

Neurodegenerative diseases are often characterized by the self-assembly of specific proteins into a fibrillar, “misfolded” amyloid state. The standard model for amyloid growth^1^ predicts that the initial nucleation is rate-limiting; the extension of fibers is assumed to be a rapid process. Recent experiments, however, have challenged this assumption. The dissolution of radiolabeled monomers from the ends of unlabeled fibers^2^ supports a two-stage binding mechanism in which the fast reversible association of monomeric Aβ to fibers (“docking”) is followed by a second, slower step, interpreted as a conformational transition of the incoming monomer on the fiber surface that results in essentially irreversible binding (“locking”).^3, 4^ Fluorescence microscopy indicates fiber elongation proceeds in a stop-and-go manner with bursts of fast elongation followed by periods of slow or absent growth^5–8^, which may be interpreted as failure of the incoming monomer to adopt the correct conformation after binding. The incorrectly docked monomer then arrests growth by preventing other incoming peptides from attaching to the fiber.^4^

A structural understanding of this process at the atomic level has proven very difficult to achieve experimentally as the growing fiber ends comprise only a small fraction of the total sample. In the absence of experimental data on the molecular level, molecular dynamics (MD) has been used to explore the dock-lock hypothesis. The fiber elongation process is too complex and occurs on too long of a time-scale to fully model at the atomic level. Coarse-grained results suggest a rugged landscape for fiber elongation^9–14^ with both productive and non-productive binding modes.^15^ On the other hand, constrained atomistic simulations starting from an geometry favorable for binding often show a smooth, downhill trajectories without prominent metastable states.^16^ Instead of the low-energy kinetic traps found in the coarse grained simulations of random coil interactions with amyloid fibers, diffusion and dehydration of the fiber tip are the primary obstacles to fiber elongation in these simulations. It is an open question how the binding of states that do not closely match the fiber conformation may initiate amyloid fiber elongation.

To explore this question for the Aβ_1-40_ peptide, we take a different approach by describing the system atomistically in explicit solvent but starting the simulation from a docked state in which the bound monomer is in a partially helical conformation reflective of some of the states in the unbound ensemble. The system eventually reaches a metastable state in which the C-terminal strand of the amyloid fiber is wrapped around a collapsed coil conformation of the monomer.

This result confirms non-productive binding modes may be a prominent feature of Aβ fiber elongation.

## Results and discussion

The simulations start from a NMR structure of the Aβ_40_ monomer in aqueous solution.17 The Aβ_40_ monomer does not have a true structure in the traditional sense, but a wealth of experimental data has shown that certain conformations are more likely than others.^18,19,20,21^ Although estimates of the actual conformations in the ensemble vary considerably, the central hydrophobic core (CHC, Leu17-Ala21) often folds into a helix in other experimental and computational studies. ^22,23,21,24^ The remainder of the peptide is largely unstructured and exposed to solvent, with the exception of the hydrophobic C- and N-ter both of which make contact with the CHC. The monomer therefore has a defined surface for docking, although the contacts defining the structure are few and weak.

The docked complex of the Aβ_1-40_ monomer with the amyloid fiber is shown in Figure 1A. Solid State NMR measurements indicate that the two β-strands of the fiber are displaced relative to each other.^25^ This stagger creates two asymmetrical shelfs at the ends of the fiber, one exposing the sidechains of the N-terminal strand and another the sidechains of the C-terminal ^s^trands. The monomer binds to the C-terminal end with the 310 helix resting on the inward facing hydrophobic residues (A22, I24, and L26). The helix is at an angle to the fiber with a slight tilt to the C-terminal strand of the fiber. The largely disordered termini of the Aβ_1-40_ monomer, which are poorly defined in the NMR structure, primarily sit above the surface of the fiber and make few contacts with the fiber surface, with the exception of R5, which makes a salt bridge to E14.

**Figure 1:**
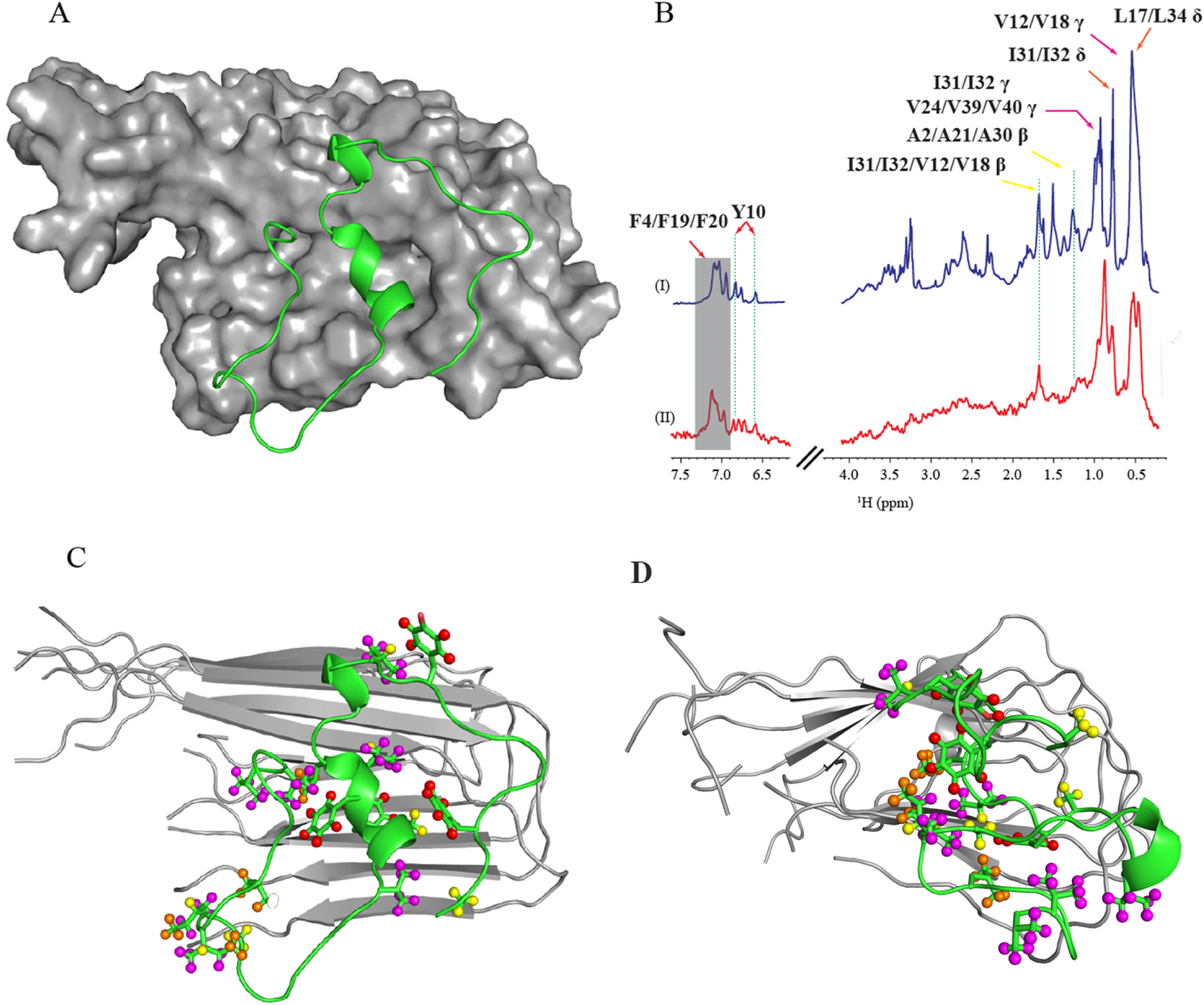
(A) The docking pose of A*β*_40_ over the surface of the amyloid fibril. (B) ^1^H STD-NMR spectrum of A*β*_40_ reflecting the group epitope mapping of the monomer to the amyloid fibril. Blue: Reference ^1^H spectra of A*β*_40_ in the presence of amyloid fibril (molar ratio of Aβ_40_ and amyloid fibril = 1:100). **Red:** STD-NMR spectra reflecting the resonance transfer to the binding epitope. (C) Comparison of the initial docked state to the STD-NMR results. The protons of A*β*_40_ in close proximity to the amyloid fibril are represented by spheres. Proton color representation: Red-aromatic proton; Yellow-beta proton; Magenta-gamma proton; Orange-delta proton. **(D)** ^D^ocked state of A*β*_40_ and amyloid fibril at the end of the simulation after 1 micro-sec MD simulation.

### Unfolding of Aβ_40_ in a stepwise transition

The docked complex was formed by rigid body docking of the experimental structured of the monomer and fiber. However, the Aβ monomer is flexible and is expected to change as it adapts itself to the surface of the fiber. Moreover, the ends of the fiber only form a small fraction of the sample and are therefore not well defined in the experimental structure. As a result, the initial complex in the simulation is not likely to be stable and will evolve as the monomer unfolds on the surface of the fiber and the ends of the fiber fray and adapt their equilibrium positions.

This transition is conveniently analyzed through cluster analysis. Clustering aims to group the conformations sampled during the trajectory to reflect the basins of the free energy landscape. Clustering requires a structural distance metric. Due to its disordered nature, the conformational ensemble is most efficiently analyzed by clustering the conformations by pair-wise RMSD. Grouping conformations by clustering offers a suitable mechanism to identify the recurrence and transition probabilities during the time course of simulation.^26^ Geometrical linkage clustering collects conformations, which are structurally similar and lie in the same energy basin of the free energy surface.

Visually, cluster analysis shows a stepwise transition away from the initial docked complex as the simulation progresses. The stepwise changes in this transition are easily tracked by changes in the inter-monomer (Figure 2) and fiber-monomer contacts (Figure S1-S3). Early points can be clustered into 5-8 closely related groups containing near-identical structures. A transition away from the initial unbound structure begins to take place within the first 100 ns. The first event is the sudden hydrophobic collapse of the protein around 50 ns, which is reflected by a decrease in both the radius of gyration (Figure S4A) and the solvent accessible area (Figure S4B). At the same time, the binding of the monomer and conformational change within a monomer, drive a corresponding structural change in the fiber. As the monomer compacts on itself, the leading C-terminal β-strand loses contact with the monomer and becomes exposed (Figures S1). The loss of the leading C-terminal β-strand drives a structural transition as the monomer and fiber change conformation to adapt themselves to each other (Figures S5). Interestingly, there is similar movement on the opposite side of the fiber. Similar to previous MD results by Okumura and Itoh,^27^ the loop connecting the C-and N-terminal β-strands at the opposite end begins to coil against itself within the first 100 ns (Figures S1-S3) mirroring the coil found between the monomer and fiber at the other end that forms as the simulation progresses (Figure 1D).

**Figure 2:**
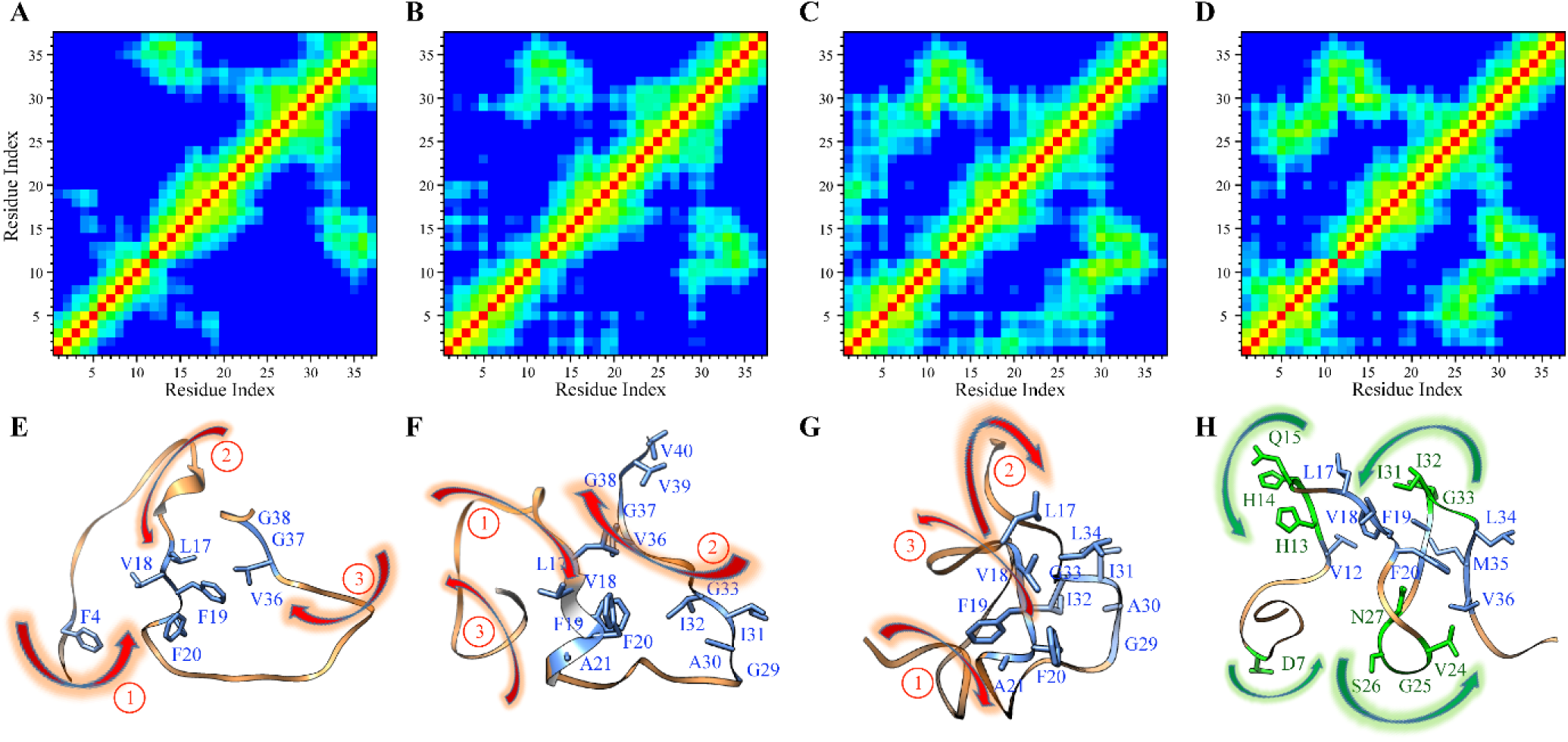
Inter-side-chain contact map (upper panel) for A*β*_40_ conformations obtained from the MD simulation in presence of amyloid fiber. **(A-D)** Contact map from time frame **(A)** 0 ns to 100 ns **(B)** 100 ns to 500 ns **(C)** 500 ns to 900 ns **(D)** 900 ns to 1000 ns from cluster centroids. The distance cut-off ranges from 0 to 1.5 Å (red to blue contours). **(E-H)** Animation of sequential phenomenon dictating the dynamics of A*β*_40_ conformations (lower panel). Sequential dynamics from **(E)** 0 ns to 100 ns. **(F)** 100 ns to 500 ns **(G)** 500 ns to 900 ns **(H)** 900 ns to 1000 ns. Sequential events are represented with red numbers and the direction of conformational change is shown with an arrow head. Key residues involved in formation of “turns” are shown in green color and residues involved in hydrophobic contact are shown in blue.

After the initial hydrophobic collapse, more contacts start to develop between the hydrophobic residues in the N- and C-terminal regions and the CHC region of the monomer, driving a structural transition to a more collapsed conformation that becomes more prominent as time goes on. During this transition, a turn forms between D23 and S26 and another between H13 and L17. Simultaneously, the N-terminus disengages from the CHC at around 500 ns. This movement is guided by a corresponding movement of the leading C-terminal β-strand to cover the part of the surface of the monomer not in contact with the fiber. The effect of these two transitions is to form a U-shaped like structure that differs from a normal anti-parallel β-hairpin by the intercalation of some of the chains between the strands. This conformation bears a slight resemblance to the repeating steric zipper found in the amyloid structure, with the exception that the secondary structure is more distorted than a canonical β-sheet and the loop connecting the β-sheets is much shorter than in the amyloid fiber. The orientation of the monomer is also inconsistent with the final “locked” conformation with the hairpin oriented nearly perpendicular to the fiber edge. Linkage cluster analysis^28, 29^ indicates this specific cluster grows at the expense of others as time progresses, eventually dominating the smaller, more structurally diverse clusters found in the 100 ns time frame (Figures S6-S7). This specific family of U-shaped structures therefore represents a distinct intermediate through which multiple conformations converge.

The system continues to evolve before eventually reaching stability around the 900 ns mark. The most prominent change in the monomer is at the C-terminal end, which bends against itself near the 900 ns mark to form another hairpin structure. This movement is matched by a corresponding movement of the leading C-terminal β-strand of the fiber into contact with the C-terminus of the monomer. Once this occurs, the system stabilizes with time. At 900 ns, the higher order clusters (4-5 groups) are of comparable size to the previous cluster and no significant shift occurs in the range of 900 ns to 1 μs time scale, suggesting near-convergence (Figure S7), at least within the limited time frame of the simulation.

### Transient metastable states found over the free energy landscape

Individual conformations are limited in their ability to describe a complex, dynamic system such as Aβ fiber binding. The analysis of the structure of cluster centers suggests the conformational transition from the bound to unbound structure is primarily driven by hydrophobic collapse. Making this assumption, energy landscapes are constructed using the radius of gyration, Rg, as one collective variable and the RMSD from the initial docked structure as the second variable to probe the conformational space within the binding process (Figure 3). The validity of an energy landscape constructed as in this manner is supported by a scree plot, which shows that 81% of the collective variance in RMSD can be explained by the first two principal components (Figure S8). To highlight the kinetic aspect of the transition, we can partition into the energy landscape into the three time regimes that were previously identified in Figure 2.

**Figure 3:**
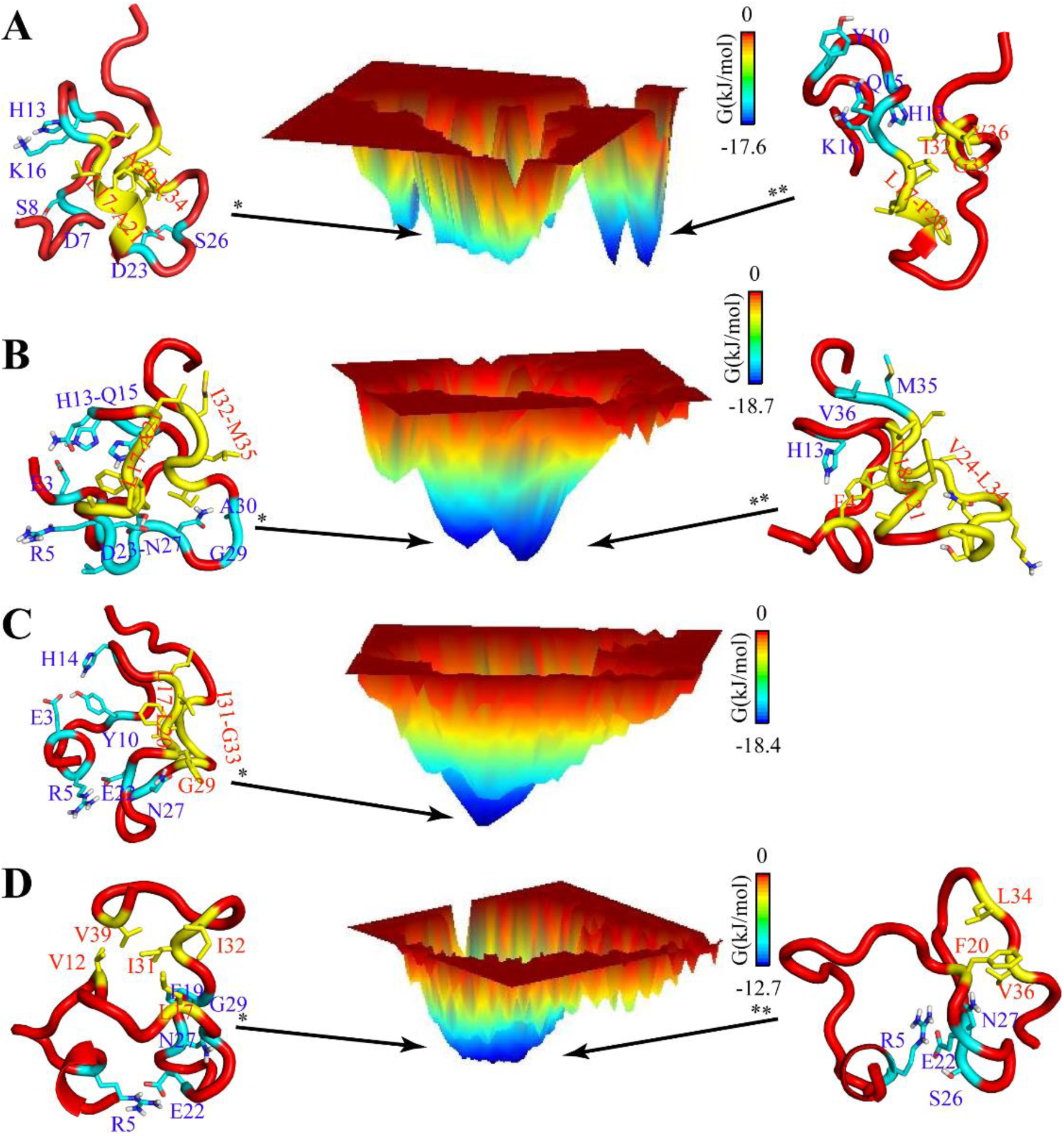
Energy landscape based on the Root-Mean-Squared Deviation (RMSD) and Radius of Gyration (Rg) as variables prepared from **(A)** 0 ns to 100 ns. **(B)** 100 ns to 500 ns **(C)** 500 ns to 900 ns. **(D)** 900 ns to 1000 ns. The Gibbs free energy cost associated with the minima structure is represented by the bar in kJ/mol. A*β*_40_ conformations representing the minima are differentiated using * and ** symbols based on the energy cost. Residues involved in formation of hydrogen bonds are represented with cyan color and those involved in formation of hydrophobic patch are represented with yellow color.

The energy landscape (Figure 3A) from the first partition has several distinct energetic minima, a broad basin with an energy of -17.6 kJ/mol closely resembling the starting structure and several sharp minima with a slightly lower free energy that correspond to more compact structures with an alternate hydrogen bonding structure. In these alternate conformations (Figure 3A left), the 3_10_ helix of the initial ^s^tructure unwinds slightly and the hydrogen bond between Asp23 and Ser26, which defines the loop that terminates the 3_10_ helix, is lost. The N-terminus also moves away from the CHC and instead folds against the N-terminal part of the 3_10_ helix. Outside the broad basin near the central minimum, the overall surface is rough, with high-energy barriers between metastable states.

In contrast to the rough energy landscape found in the initial phases of the simulation, the energy landscape from the 100-500 ns time is smooth with two deep but broad minima separated by a low energy barrier, one dominated by a series of bend structures (Figure 3B right) and another dominated by the hydrophobic collapse of the C-terminus against the CHC (Figure 3B left). A conformational change in the loop region from V24-I32 converts either of these structures into the minimum energy conformations found between 500-1000 ns. The conformational change is driven by a change in the hydrogen-bonding network of the polar side chain of Asn27.^22^ In the earlier structures, the Asn27 side chain is hydrogen bonded to the adjacent backbone residues. After the appearance of dominant cluster at ∼500 ns (as discussed in Figure S7), the side chain moves in between the two strands of the loop, creating a hydrogen bond network between the backbone amides of Asn27 and Phe19; and Glu22 on one strand of the loop and Gly29 on the other. The side chain of Asn27 therefore appears to act as a switch to converting one conformation to another. Importantly, the hinge region at Gly25-Asn27 is well-maintained in all the structural ensembles, which has been reported as key conformational change promoting aggregation in several previous reports.^30, 31^ Turns and bending within the regions Glu7-Gly9, His13-Gln15, and Val24-Asp27 also correlate well with earlier findings.^32,33,34,35^ The transition to final conformation is largely mediated by a change on the fiber side. The hydrophobic Val24-Ile31 loop, which had previously become disordered with the loss of the final β-strand, at around 700 ns forms a structured loop that packs against the end of the pseudo-hairpin of the monomer. The final state is at the bottom of a broad energetic minimum that represents a large ensemble of closely related structures, reflecting the fact that convergence has been reached.

### Validation of the converged structure with STD-NMR

The accuracy of the docked model was checked by STD NMR, which detects transient ^c^ontacts between the Aβ_1-40_ monomer and high molecular weight species by spin diffusion (Figure 1). The interpretation of the spectrum was complicated by line broadening from field inhomogeneity caused by the large fiber particles resulting in severe signal overlap. Due to the degree of signal overlap, peaks could not be uniquely assigned and a quantitative analysis is impossible. Nevertheless, even with the overlap, a qualitative estimation of the relative strengths of interactions among different types of residues could be made.

The strongest STD signal arises from the terminal protons from branched aliphatic residues: γ protons from Val12 and Val18, δ protons from Leu17 and Leu34, suggesting that these hydrophobic residues are most frequently in contact with the fiber and likely contribute most of the bulk of the binding energy (Figure 1B). The aromatic rings of the Phe19 and Phe20 residues also make frequent contacts with the fiber surface. Both findings are reflected in the contact map of the MD simulation (Figure S6), at least on a qualitative level. Specifically, F20 makes multiple contacts with the fiber. The terminal δ and γ of Ile31 and Ile32 and the β protons of Ala21 and Ala30 show a somewhat weaker STD signal. The STD effect is noticeably weaker in the Hα region 4.0-4.5 ppm. The lack of an STD signal in the Hα region indicates most of the contacts of the fiber are to the side chain of the A*β*_40_ monomer and not the backbone (Figure 1C and 1D), in agreement with previous CPMG data.^36^

### Estimates of Markov States

Based on the conformational analysis of the trajectory, convergence is apparently achieved at ∼900 ns from a transition from a metastable state occurring at around 400 ns. The apparent convergence does not mean that the system is static merely that the energy landscape is no longer changing on the nanosecond to microsecond time-scale. Complex biological systems often have multiple energy wells separated by high barriers that are inaccessible to molecular dynamics simulations with conventional sampling. The transition from the initial docked state to the final locked conformation takes place on a time-scale of tens of seconds, implying a series of rare events are evolved that are not sampled with a 1 µs molecular dynamics simulation.

To further explore the conformational space at convergence, we first performed accelerated molecular dynamics (aMD) to starting from ten conformations taken from snapshots at equal intervals within the 400 to 900 ns time segment. This method is an enhanced sampling technique that reduces the energy barrier that separate the states in the conformational hyperspace to favor transitions between energy minima.^37, 38^ A dual boosting scheme using the entire potential and a separate boosting term to the torsional energy terms of the peptide specifically was employed to accelerate both large scale movements of the protein and solvent (Table 1).

**Table 1:** The details of the aMD parameters used in the simulation. These values are obtained from a preparatory 10 ns classical MD for each of the system.

| aMD Systems | Total number of atoms | Average Potential Energy (kJ/mol) | Dihedral Energy (kJ/mol) | EthreshP | alphaP | EthreshD | alphaD |
| --- | --- | --- | --- | --- | --- | --- | --- |
| 1 | 28855 | -85817.751 | 2273.009 | -81201.0 | 4616.8 | 3201.009 | 185.6 |
| 2 | 26104 | -77101.553 | 2249.983 | -72924.9 | 4176.64 | 3177.983 | 185.6 |
| 3 | 27271 | -80831.570 | 2246.226 | -76468.2 | 4363.36 | 3174.226 | 185.6 |
| 4 | 27661 | -82075.817 | 2255.547 | -77650.1 | 4425.76 | 3183.547 | 185.6 |
| 5 | 31570 | -94518.129 | 2248.865 | -89466.9 | 5051.20 | 3176.865 | 185.6 |
| 6 | 29566 | -88147.302 | 2234.292 | -83416.7 | 4730.56 | 3162.292 | 185.6 |
| 7 | 27388 | -81218.981 | 2250.990 | -76836.9 | 4382.08 | 3178.990 | 185.6 |
| 8 | 26767 | -79192.512 | 2251.883 | -74909.8 | 4282.72 | 3179.883 | 185.6 |
| 9 | 26707 | -79013.412 | 2252.871 | -74740.3 | 4273.12 | 3180.871 | 185.6 |
| 10 | 26611 | -76072.783 | 2257.088 | -71815.0 | 4257.76 | 3185.088 | 185.6 |

A network model mapping the transitions between states can be constructed from these 10 short trajectories. A meaningful network model can only be created if the many conformations from the simulations are first clustered into a small number of states. Directly using atomic distances as variables for clustering produces a sparsely populated high dimensional space that leads to poorly defined clusters. The dimensionality of this space can be reduced by using principal component analysis but clustering based distances can give improper weight to the large amplitude fluctuations typical of unstructured regions, which are largely irrelevant for distinguishing between important conformational states.

An alternative is to consider the kinetics associated with each state, grouping conformations that undergo similar large-scale motions rather than similar conformations by the RMSD definition. Kinetic variables that serve as a reaction coordinates can be defined based on the eigenvalue decomposition of a time lagged correlation matrix (time-lagged independent component analysis (TICA)). The time lag defines the time-scale of motion; shorter time lags correspond to high frequency motions while longer time scales correspond to more rare collective events. A plot of lag time versus the implied relaxation time scale suggests a time-lag of 3 ns, adequately models the slow collective motions in the sample (Figure S8). A Hidden Markov Model network can then be constructed after clustering the states based on the kinetic variables identified by TICA.

### Network transition pathway analysis

The five macrostates generated by fuzzy clustering are easily identifiable as basins on an energy map (Figure 4) that can explain 85% of the kinetic variance (Figure 5 right). State 0 is a collapsed coil conformation with a pseudo hairpin between that closely resembles the final, converged conformation obtained from cluster analysis (Figure 2H). States 1 and 2 are both anti-parallel β-sheet hairpins with a short β-strand from Y10-V12. The two states are differentiated by the other strand in the β-hairpins pair: in State 1 the Y10-V12 β-sheet is paired with E3-H6 near the N-terminus while State 2 is paired with a C-terminal strand from L34-V36. This is a different conformation then the anti-parallel β-sheets found in D23N mutants of Aβ where the C-terminal β-sheet is paired with another C-terminal β-sheet reflected about the fiber axis.^39^ State 3 is a more collapsed version of state 0 generated by the wrapping of the C-terminus against the CHC is stabilized by van der Waals and electrostatic forces.^18^ State 4 is a mostly unfolded extended conformation characterized by bends and devoid of other secondary structure. State 4 is also closest to the amyloid conformation, as it can be converted into the fiber conformations by several crankshaft motions and the straightening out of the β-bends into two regular β-sheets. The remaining conformations require the breaking of secondary structure and tertiary contacts.

**Figure 4:**
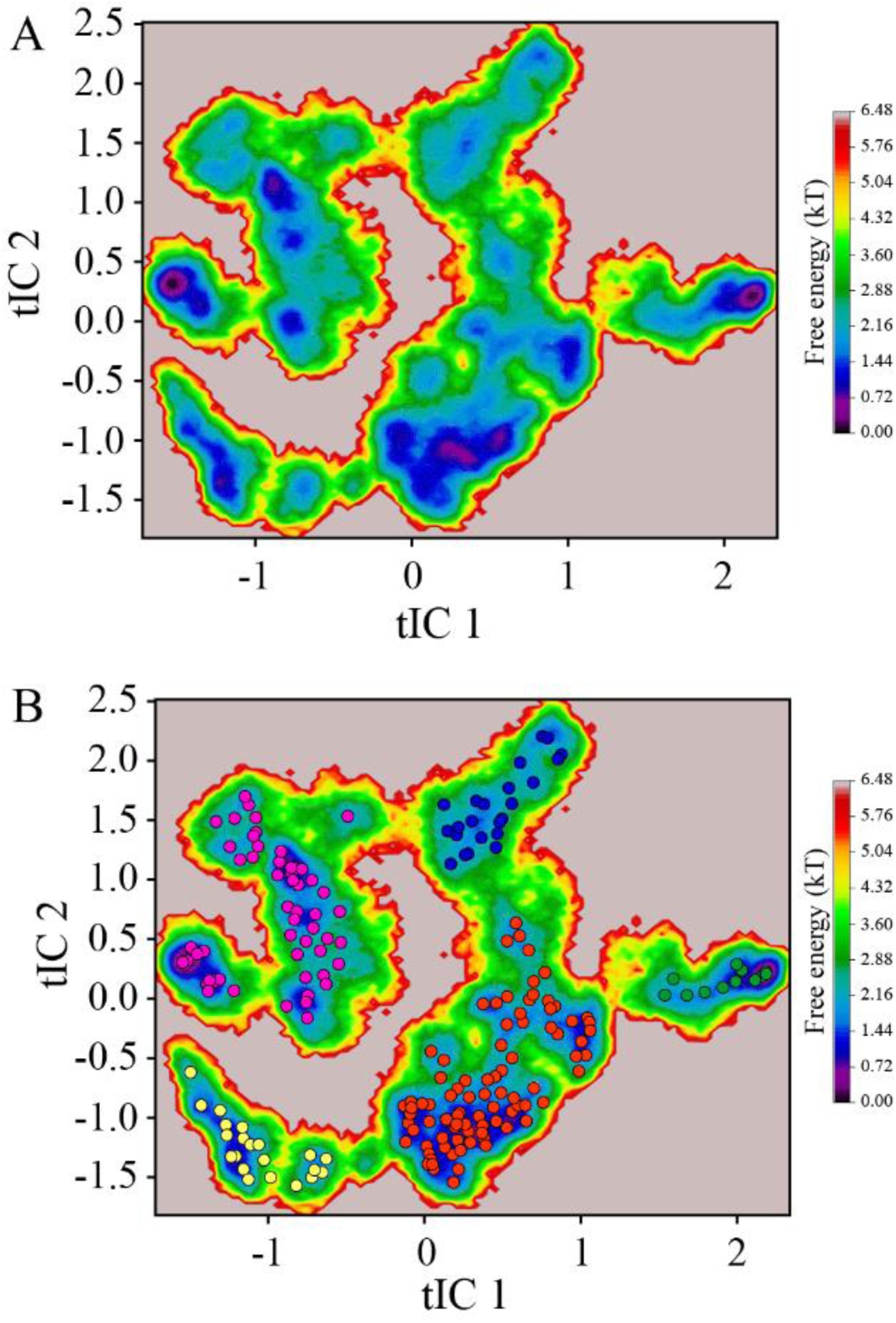
(A) Free energy map of the Hidden Markov State Model plotted against tICA component 1 and 2. (B) Projection of the 200 microstates onto the free energy map. The microstates are lumped into 5 macrostates, which are colored according to their membership.

**Figure 5:**
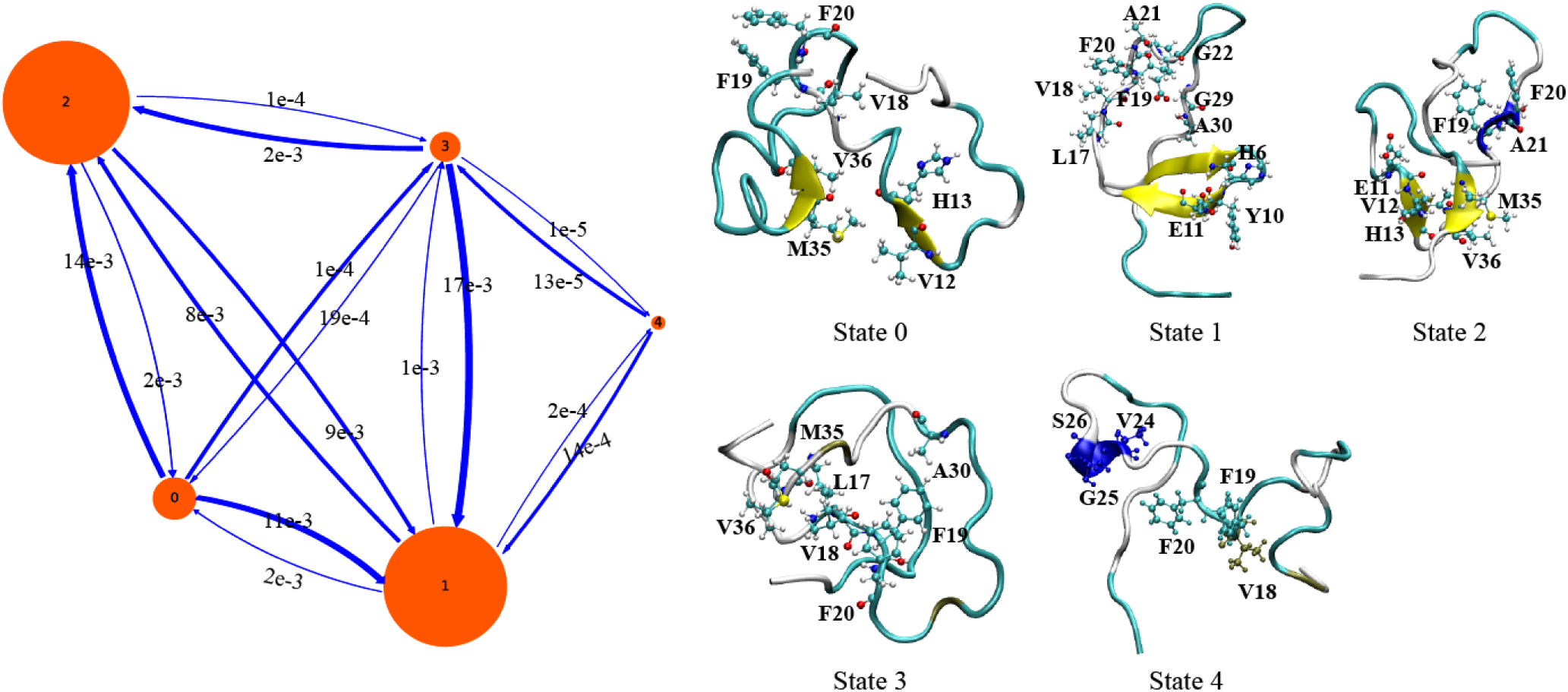
Network transition pathway analysis of the Hidden Markov State Model. The probability flux between the 5 macrostates is shown on the left, with the arrow size proportional to the probability of transition. Representative conformations of the 5 macrostates are shown on the right with key residues outlined with the CPK model.

A network transition pathway between the states can be constructed using the Hidden Markov Model. State 0 is near the minima of the energy landscape at the end of the simulation and can be thought of as the starting state of the model. From this starting state, the transition flux is outward, primarily to states 1 and 2, with some flux towards state 3. States 1 and 2 interconvert with each other but transitions to other species are rare. State 3 serves as an intermediate between State 0 and States 1 and 2 with most of the flux coming inward from State 0 and outward to States 1 and 2. State 4 is essentially a rare off-pathway intermediate along the transition from State 3 to State 1. The barrier between states is relatively low, averaging around ∼2-3 kT (1.2-1.8 kcal/mol). Overall, the model suggests a transition after binding from a partial helical conformation to a series of collapsed coil structures into beta-sheet hairpins that interconvert on a slower time-scale.

## Discussion

The transition of the monomer from the initial bound state to the final metastable intermediate appears to be the consequence of five distinct sequential events: 1) Unfolding of the 3_10_ helix on the surface of the fiber. 2) Hydrophobic collapse and compaction following the expulsion of water from the interior. 3) Movement of the leading C-terminal β-strand of the fiber away from the main body of the fiber and into contact with the monomer. 4) A concerted movement within the monomer involving disengagement of the N-terminus away from the central hydrophobic cluster and the creation of two hydrogen bonded turns to form a distorted hairpin structure stabilized by the interaction of sidechains of the β-strands against each other rather than by hydrogen bonds between strands (Figures 2H, 3D, and state 0 in Figure 5). 5) Packing of the leading loop of the pseudo hairpin against the monomer surface to generate the first metastable state. 6) Further evolution to form two interconverting β-hairpin stabilized by hydrogen bonds (Figure 5 - States 1 and 2). Overall, the transition can be divided into two distinct steps: a fast internal unfolding and compaction of the monomer and slower rearrangement driven by changes in the fiber conformation.

The final product of the simulation is a network model of the conformational rearrangements occurring after the unfolding of the monomer on the surface of the fiber. Two anti-parallel β-sheet conformations dominate the conformational landscape. Neither conformation is compatible with the conformation of the amyloid fiber, implying a further conformational transition is needed for fiber extension. On the basis of surface plasmon resonance experiments, this step occurs in a series of events predicted to occur on the seconds time-scale,^40^ beyond on the time-scale of our simulation experiments. The conformational states in the network model therefore likely correspond to the initial “docked” states in the dock-lock hypothesis. Most of the conformations are incompatible with incorporation into the fiber and are kinetic traps that must be overcome for fiber extension. One relatively rare state (Figure 5 - state 4) can likely be converted into the amyloid conformation by a series of crankshaft motions and may represent a possible pathway out of the kinetically trapped docked conformations and into a growth competent conformation.

Any simulation necessarily requires a choice between conformational sampling and physical accuracy. To reduce the degrees of freedom to a tractable number and enable specific water and interactions, we started the simulation from a docked complex using an experimental structure as the initial basis for the monomer conformation. The accuracy of the simulation therefore depends on the plausibility of the initial model. The defining characteristic of our initial starting structure is the 3_10_ helix in the central hydrophobic core of the monomer. Helical structures have been observed as both elements of the conformational ensemble of the Aβ_1-40_ /Aβ_1-42_ monomer^41–44^ and as discrete oligomeric intermediates along the aggregation pathway.^45–50^ Although the exact details of the surface induced unfolding process will likely vary for different initial starting conformations, the basic elements of the conformational transition; namely the loss of non-β-sheet secondary structure, the compaction of the monomer following expulsion of water from the interior, and the envelopment of the exposed hydrophobic elements in the monomer by the frayed ends of the fiber, are likely conserved. The simulation here may serve as a model for the binding of other non-β-sheet conformations to amyloid fibers, both as a step in fiber elongation and in the entrapment of other non-amyloid proteins at the fiber ends,^51, 52^ which may play a vital role in amyloid cytotoxicity then by preventing them from carrying out their biological roles.^53, 54^

## Materials and Methods

All the simulations were carried out using the Amber14 suite with the ff99SB-ILDN^1^ force-field on a GPU workstation having a C600/X series Tesla card. Initial coordinates for the molecular modelling study were collected from the Protein Data Bank, accession codes 2LFM and 2M4J, for the partial helical17 and amyloid fiber structures55 of Aβ_1-40_ (amino-acid sequence DAEFRHDSGYEVHHQKLVFFAEDVGSNKGAIIGLMVGGVV), respectively. The initial docked conformation of monomeric Aβ_1-40_ complexed with amyloid fiber was constructed using the Z-dock server,^56^ which uses a rigid body protein-protein docking algorithm based on the Fast Fourier Transform correlation technique to explore the translational and rotational space about the complex. The selection of docked poses within clusters are based on the pairwise-shape complementarity and energies of atomic contact, desolvation, and electrostatic parameters.^56^ The final selection of docked complex in this work is based on the visual inspection as well as the low Z-Rank-score.^57^

### Classical MD simulation

We performed an unbiased simulation procedure within the uniform density approximation with application of periodic boundary conditions (PBC) in the three-dimensional space. The simulation began with solvation of the complex with TIP3P water models in a truncated octahedron water-box with an edge-length of 10 Å from edge solute atom.^58^ Neutralization of the solvent system was achieved by addition 7 Na^+^ counter ions added using a Monte-Carlo simulation method.^59^ 10,717 water molecules were added for a total of 35,620 atoms. The length of the periodic box is 8 Å from the edge atoms of the docked complex. The PME method with cubic *β*-spline interpolation and a grid spacing of 1 Å was used for calculation of the electrostatic interactions.^60^ For the computation of the Lennard-Jones potential and Coulomb interactions, the non-bonded cut-off was fixed at 1.0 nm. The SHAKE algorithm was used for re-ordering the hydrogen bond length during simulation with an integration time step of 2 fs and a relative tolerance value of 10^-5^ Å.^9^

Energy minimization was first performed to relax the water molecules, holding the protein and ions of the system constant, for 2000 steps using the steepest descent algorithm. Following this, the protein system was relaxed, restraining the water molecules for equivalent steps. Finally, the entire system was minimized with 2000 steps using the steepest descent algorithm, followed by conjugate gradient minimization until the RMS fluctuations and gradient were stable. The system was then heated to bring the temperature to 300 K using Langevin dynamics.^61^ During the uniform density approximation, constant pressure dynamics were performed with isotropic position scaling using the Berendsen barostat to maintain correct density.^62^ The temperature in the production run was controlled using the canonical (constant T) ensemble. The total production run was 1000 ns (1 µs). The analysis of trajectory frames was performed using tools MMTSB for cluster analysis,^12^ PCAsuite for principal component analysis (http://mmb.pcb.ub.edu/software/pcasuite/), cpptraj^13^, Bio3d,^14^ VMD for visualization and timeline analysis,^15^ and various GROMACS modules^63^.

### Accelerated MD simulation (aMD)

Ten conformations taken from snapshots at equal intervals within the 400 to 900 ns time segment were subjected to a 10 ns classical MD simulation to obtain the parameters for a 50ns accelerated MD simulation to improve sampling by decreasing the energy barrier between high-energy states.^38^ A dual boosting system^37^ was used using the dihedral potential and the total potential of the system as independent boosting energies defined according to

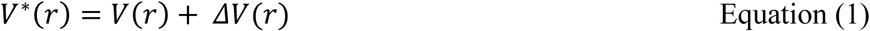

***V*(r)*** in equation 1 refers to the modified potential and ***ΔV(r)*** refers to the boost potential. The boost potential was defined by equation 2.^64^

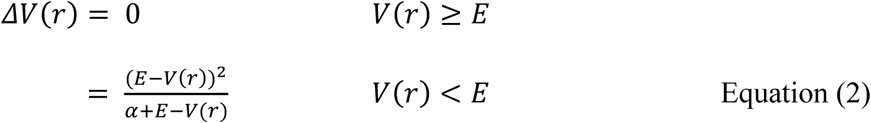

where ***E*** refers to the threshold energy, which governs the potential surface affected by the boost, whereas ***α*** is the acceleration factor determining the shape of the modified potential. Values of ***E*** and ***α*** are calculated from number of atoms and the total energy from a 10 ns preparatory simulation and can be found in Table 1.

### Trajectory Analysis

The Hidden Markov State Models (HMSM)^29^ were constructed using PyEMMA *2.4*^65^ and the Jupyter notebook.^66^ based on the cosine values of the backbone torsion angles (phi *Φ* and psi *Ψ*), and the distances between the C*α* atoms of Leu17, Val18, Gly29, and Val36 as discretization features. This distance criteria was used to include the interaction of LVFF region with the C-ter residues that have active role in governing the amyloid-beta conformations.^20^ These distances were converted into kinetic variables using time-lagged independent component analysis (TICA).^67^ The *k-means* clustering algorithm^68^ was further used for categorizing the conformation into 200 microstates. The PCCA+ algorithm was finally used to cluster the kinetically relevant microstates into 5 final macrostates (or statistically relevant meta-stable states).^69^

The transition process can only be considered Markovian if the probability for future states,^70^ can be predicted for present state according to Equation 3.

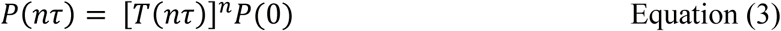

In which ***P*** is the vector of the state populations, ***τ*** is the lag time, and ***T*** is transition probability matrix, of the model. The Chapman-Kolmogorov test^71^ was used to validate the Markovian (Equation 5).

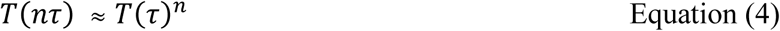

Where, **T(τ)** is an estimate of transition matrix with reference to the lag time ***τ*** and ***n*** is the number of integration steps. The results of the test are shown in Figure S9. This test is performed with the help of *cktest* script.^65^ The free energy is constructed with Equation 5:

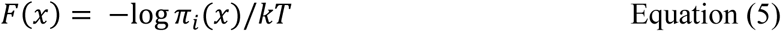

In Equation 5, π_*i*_ refers to the stationary probability of state ***x*** estimated with reference to the first eigenvector of the HMSM transition matrix.

### Saturation Transfer Difference (STD) NMR spectroscopy

Saturation transfer difference (STD) NMR experiments were performed on a Bruker AVANCE III 500 MHz NMR spectrometer with a 5 mm SMART probe at 288 K. Unlabelled Aβ_1-40_ was purchased from Genscript USA, Inc. To remove preformed aggregates, the peptide was dissolved in 1 % ammonium hydroxide at 1 mg/ml followed by the removal of the solvent by lyophilization. After lyophilization, 0.1 mg of the peptide was first dissolved in 270 µL of 1 mM NaOH in 10% D_2_O at 4°C and sonicated for 15 min. 30 µL of 200 mM sodium phosphate (pH 7.2) was added to bring the final concentration to 78 µM in 20 mM sodium phosphate (pH 7.2). The sample solution was finally brought to room temperature and filtered through a 0.22 μm filter immediately before the start of each experiment. For preparation of the Aβ_1-40_ fibers, 0.1 mg of the lyophilized peptide was solubilized in a 10 mM sodium phosphate buffer (pH 7.4) solution at a concentration of 160 μM and incubated for 48 hours at 37 °C under agitation at 1000 rpm. ∼1 μM mature cross-β fibril was added to 78 μM of the monomeric Aβ_1-40_ for the STD experiment.

## Acknowledgements

This research was partly supported by Institutional fund (Plan Project-II to A.B.), and partly by Council of Scientific and Industrial Research (CSIR), Govt. of India (02(0292)/17/EMR-II) (to AB).

## Supplementary Information

**Figure S1:**
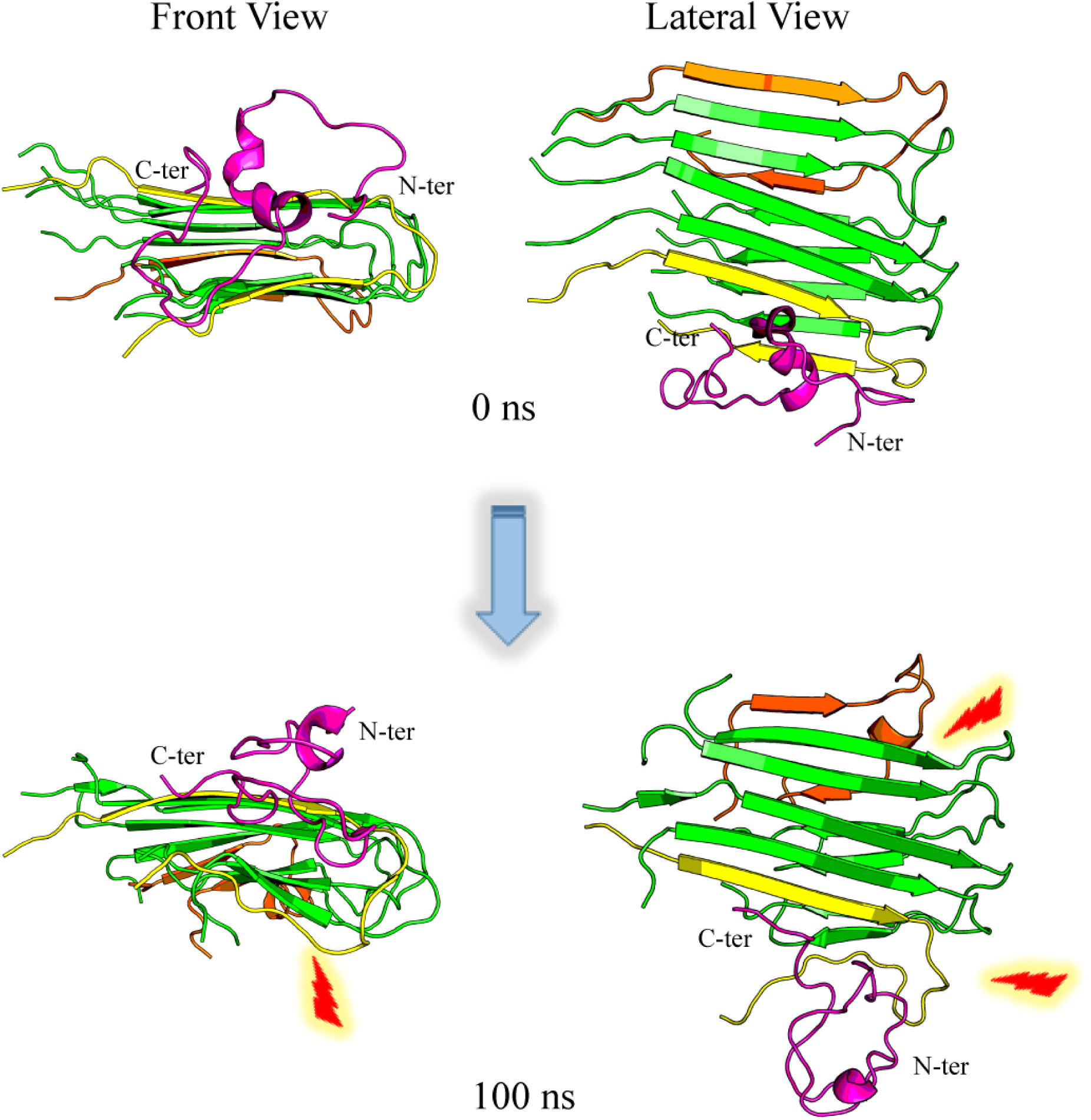
Step-wise changes: Fiber-monomer contacts. Analysis of the molecular fluctuations in the amyloid fiber in the presence of Aβ_1-40_ from 0 ns to 100 ns. The incoming Aβ_1-40_ monomer docked to the amyloid fiber is shown in pink color. The fragment to which Aβ_1-40_ is docked (near-fragment) is shown in yellow and the extreme end (far-fragment) is shown in orange. Significant changes in the secondary structure of near and far-fragment are shown with red highlights.

**Figure S2:**
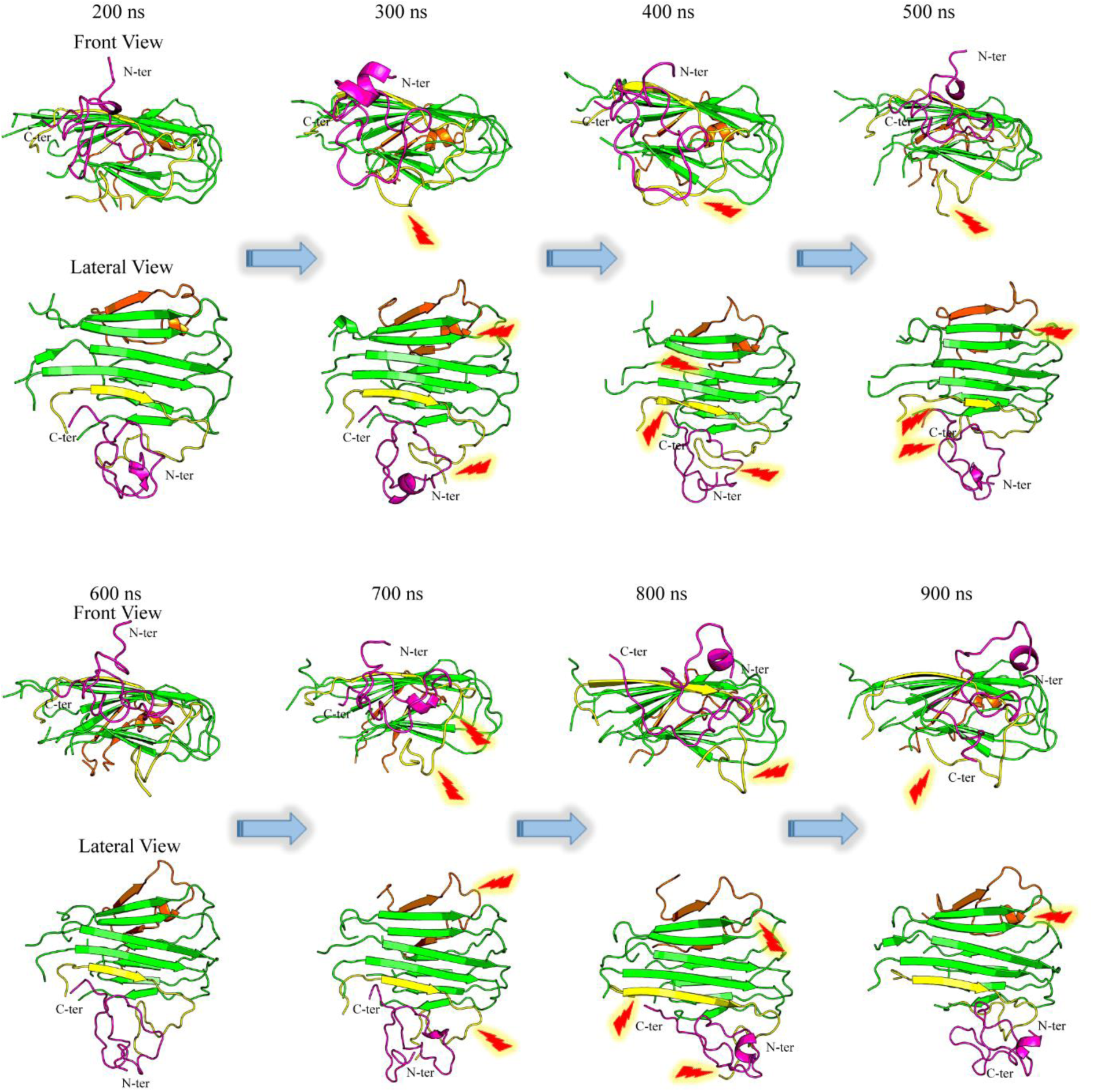
Analysis of the molecular fluctuations in the amyloid fibrillar structure in presence of Aβ_1-40_ from 200 ns to 900 ns. The incoming Aβ_1-40_ monomer docked to the amyloid fiber is shown in pink color. The fragment to which Aβ_1-40_ is docked (near-fragment) is shown in yellow and the extreme end (far-fragment) is shown in orange. Significant changes in the secondary structure of near and far-fragment are shown with red highlights.

**Figure S3:**
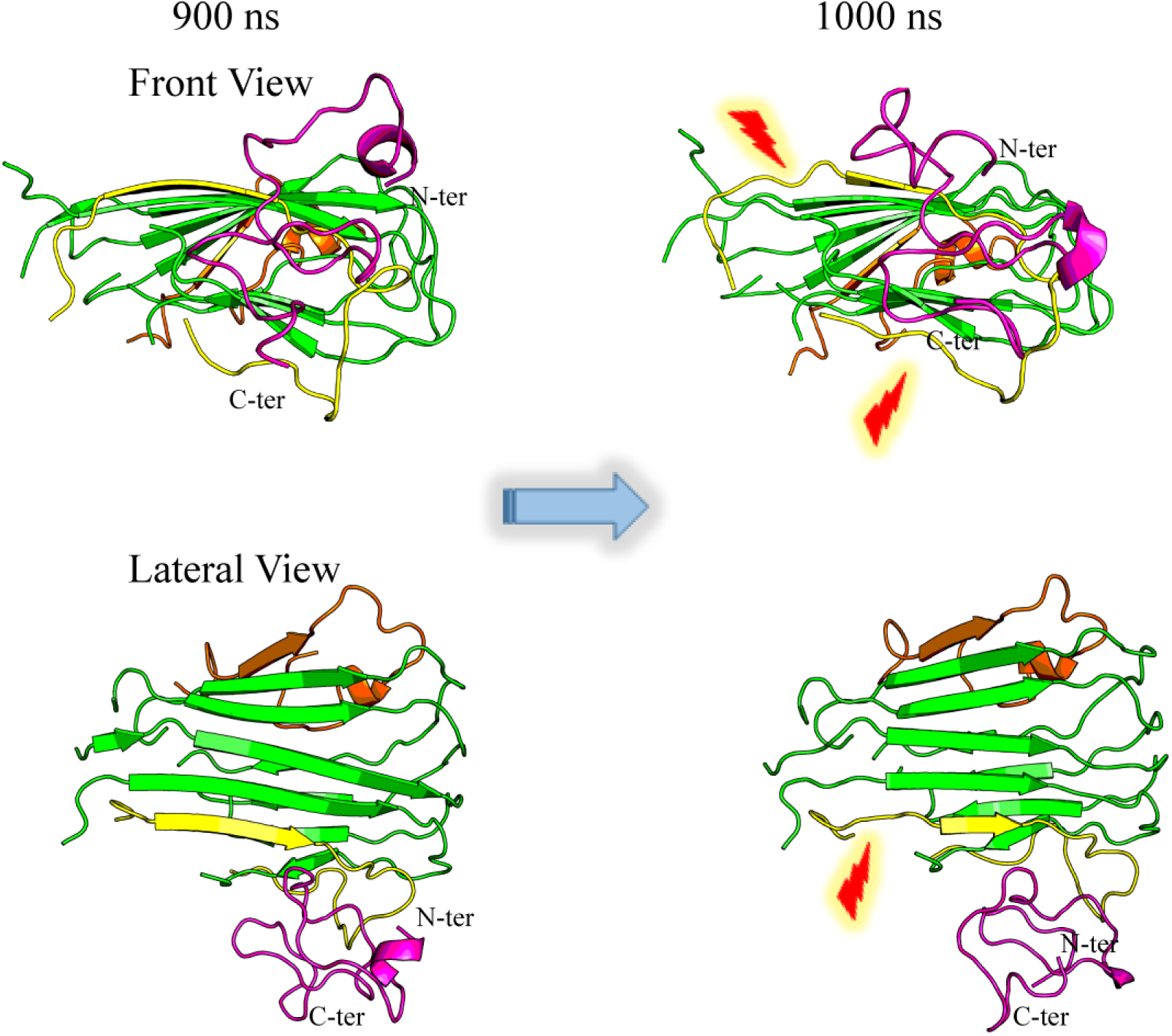
Analysis of the molecular fluctuations in the amyloid fibrillar structure in presence of Aβ_1-40_ from 900 ns to 1000 ns. The incoming Aβ_1-40_ monomer docked to the amyloid fiber is shown in pink color. The fragment to which Aβ_1-40_ is docked (near-fragment) is shown in yellow and the extreme end (far-fragment) is shown in orange. Significant changes in the secondary structure of near and far-fragment are shown with red highlights.

**Figure S4:**
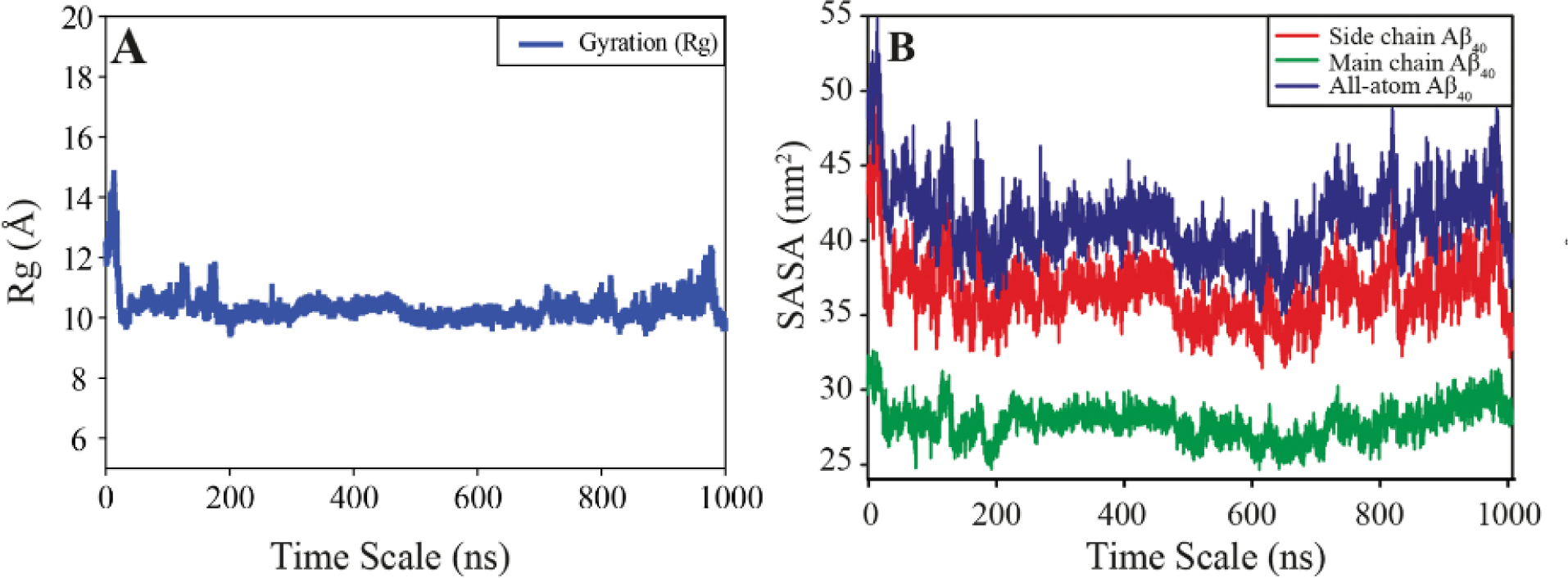
(A) Plot of the radius of gyration (Rg) of the monomer from the MD simulation. (B) Plot of the change in solvent accessible surface area as a function of time

**Figure S5:**
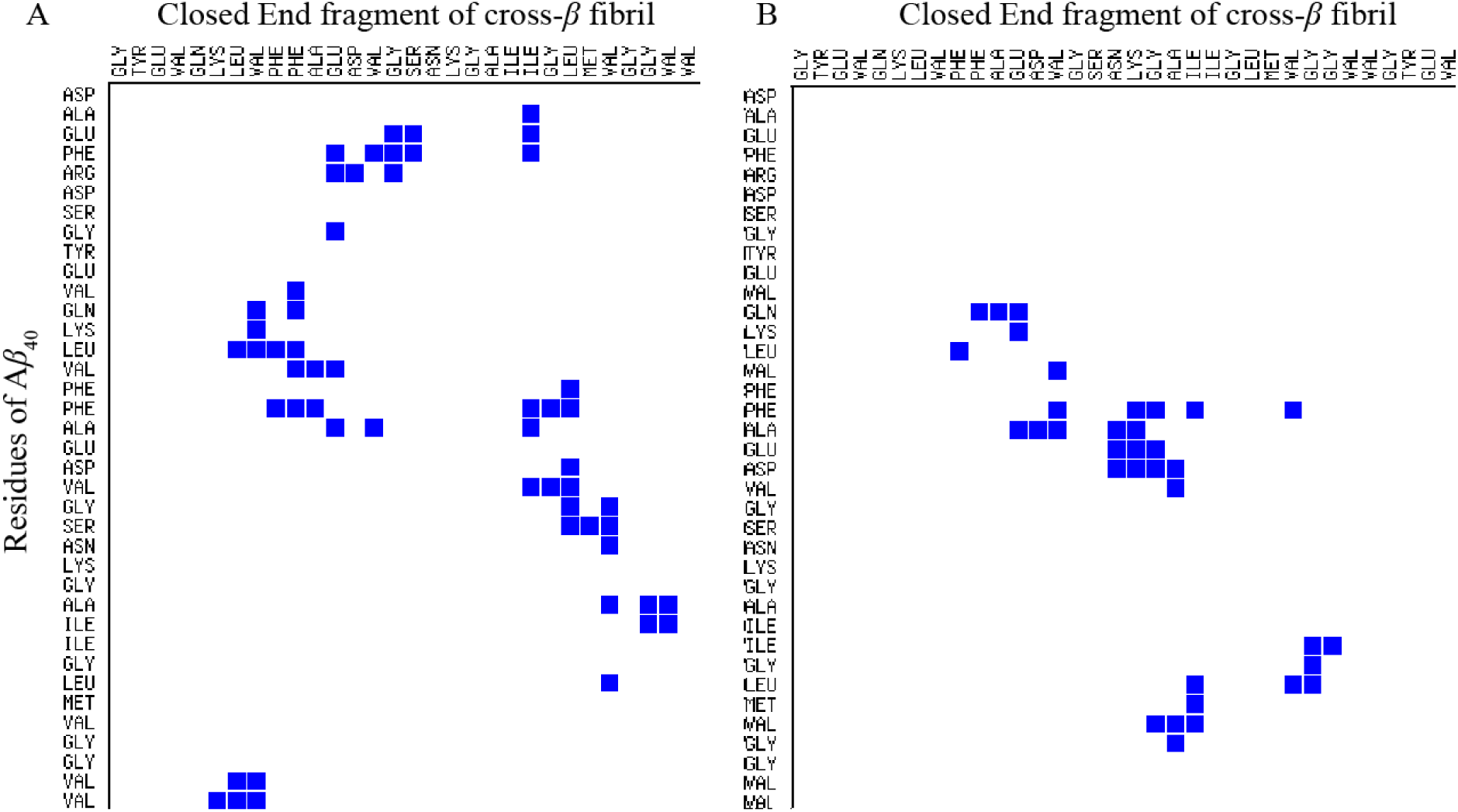
Contact map analysis between the fiber and monomer of complexed Aβ_1-40_ at (A) the start of the simulation (0 ns) and **(B)** end of the simulation (1000 ns). The plot was computed using Contact Map Analysis server (ligin.weizmann.ac.il/cma/).

**Figure S6:**
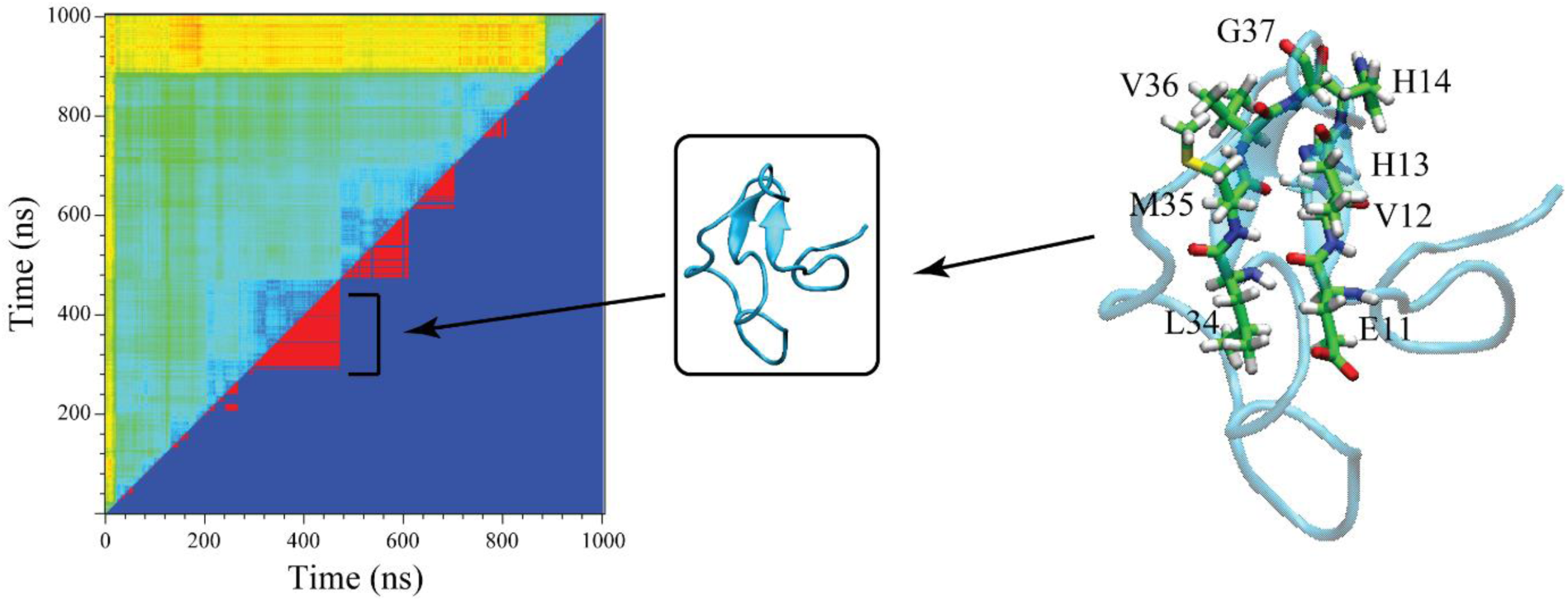
Representative structure from the largest cluster which appears at ∼500 ns. Residues that represent *β*-sheet secondary structure are shown in extreme right panel represented as sticks.

**Figure S7:**
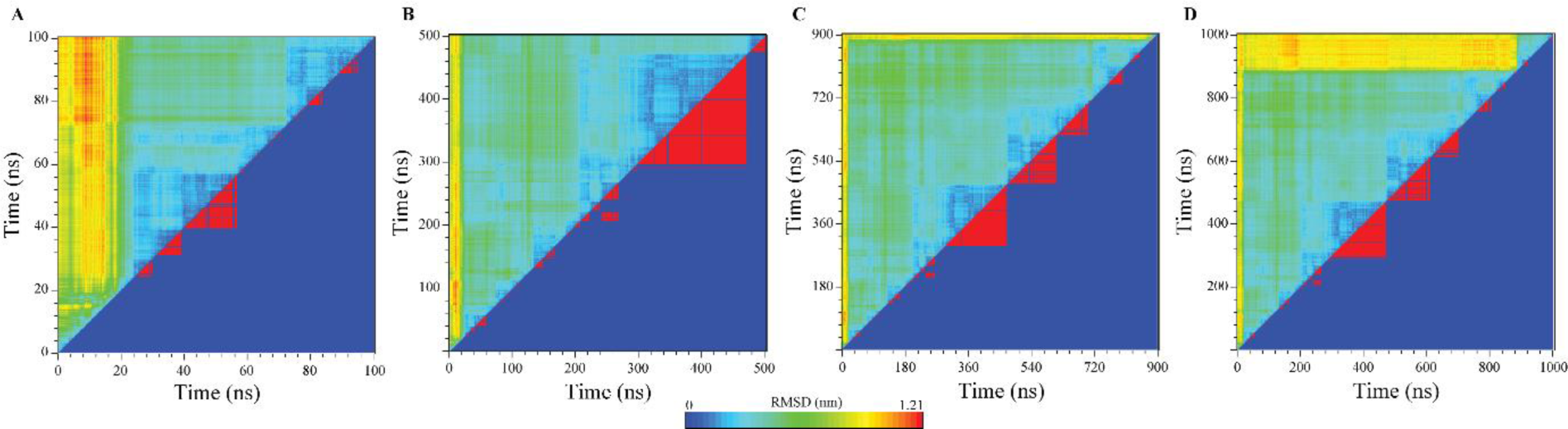
Cluster index based on pairwise RMSD plot for Aβ_1-40_ conformations, analysed for (A) 0 ns – 100 ns, **(B)** 500 ns, **(C)** 900 ns, **(D)** 1000 ns. The RMSD cut-off ranges from 0 to 1.21 Å (red to blue contours).

**Figure S8:**
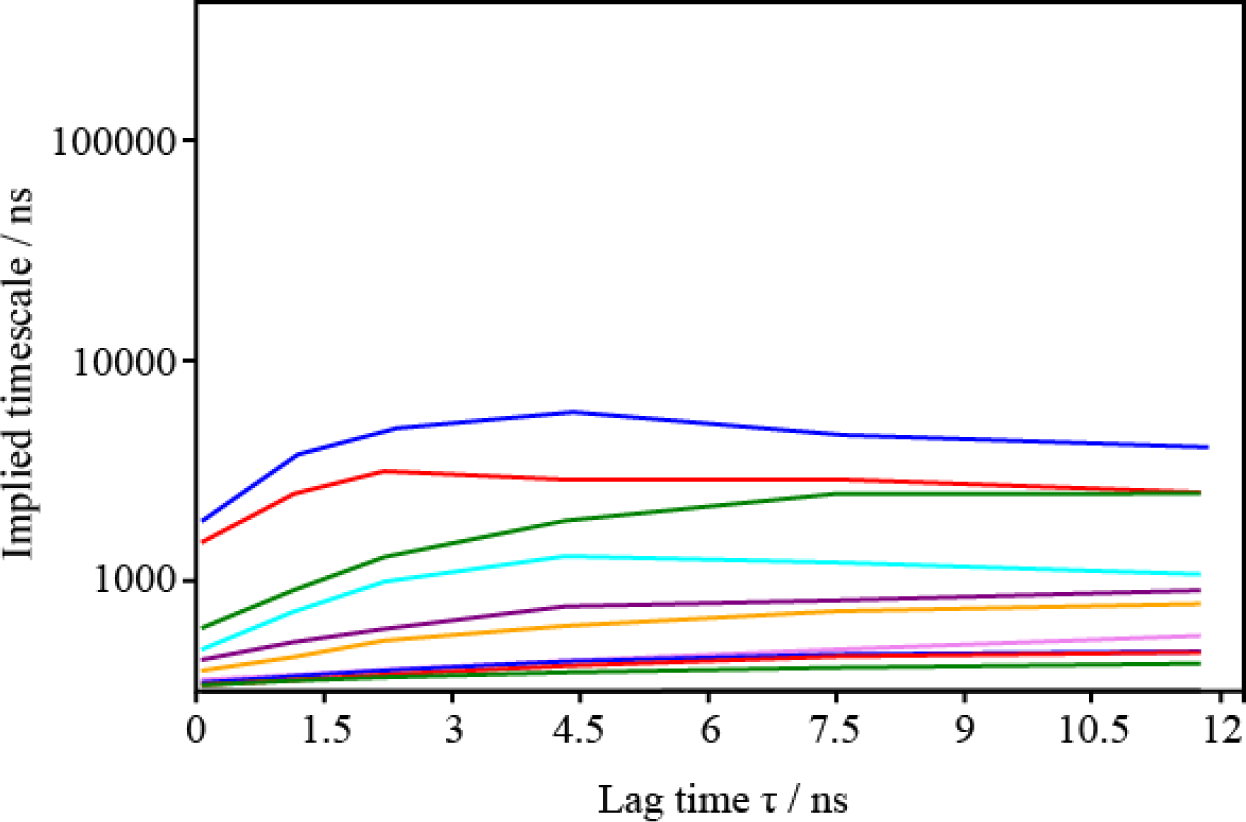
Implied relaxation time scale of the MSM constructed by the *k*-means algorithm constructed with 200 centers.

**Figure S9:**
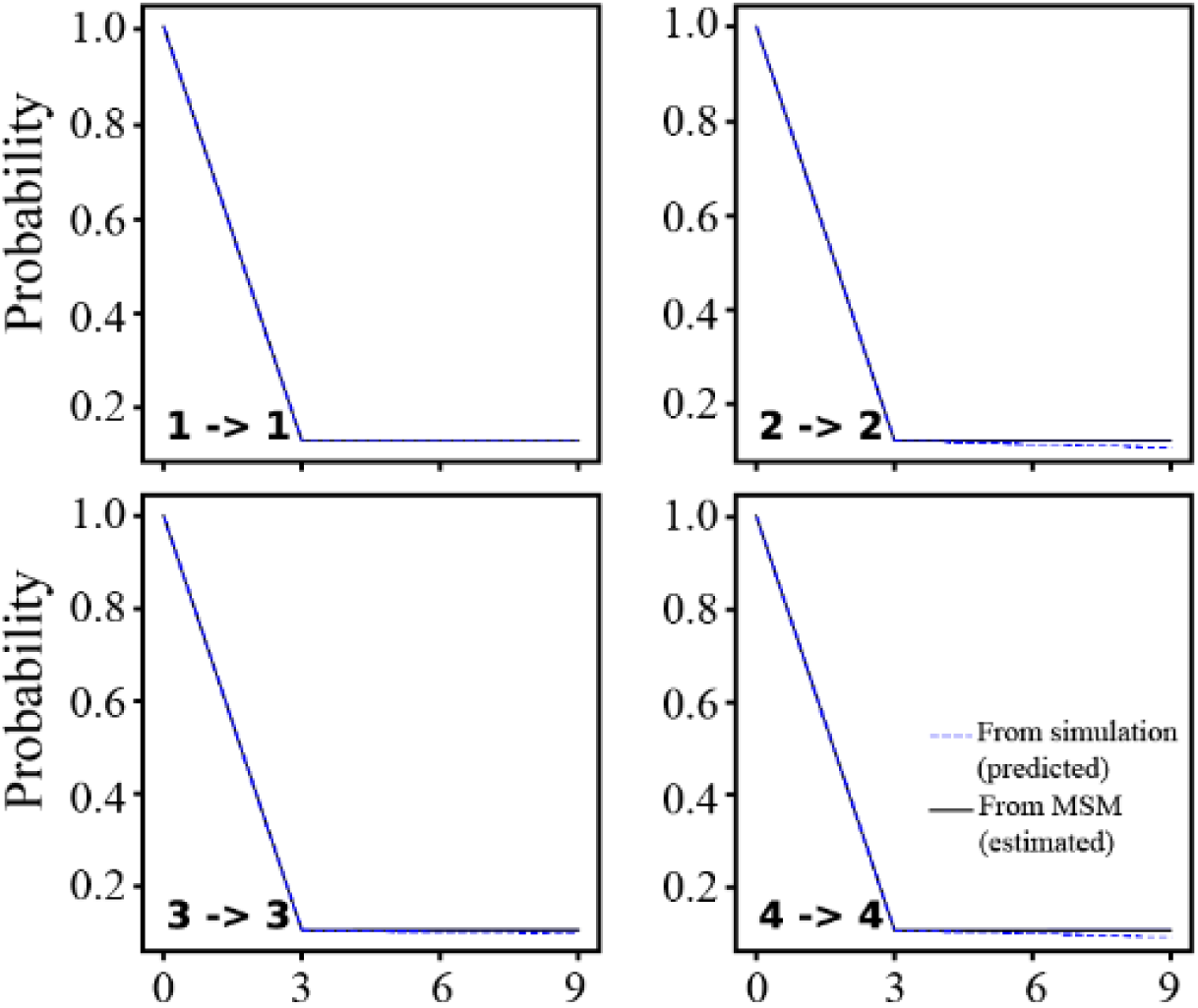
Results of the Chapman-Kolmogorov tests for the MSM at lag τ = 3ns.

